# The genetics of fruit skin separation in date palm

**DOI:** 10.1101/2024.07.01.601480

**Authors:** Shameem Younuskunju, Yasmin A. Mohamoud, Lisa Sara Mathew, Klaus F. X. Mayer, Karsten Suhre, Joel A. Malek

## Abstract

The physical appearance of date palm (Phoenix dactylifera) fruit (dates) is important for its market value. Many date-producing countries experience significant financial losses due to the poor appearance of the fruit, skin separation or puffiness being a major reason. Previous research showed evidence linking the skin separation phenotype to environmental conditions. In this study, we show that there is both an environmental and genetic contribution to the fruit skin separation phenotype. We show that beyond environmental factors, genetics is a strong contributor to the most extreme skin separation in some cultivars. A genome-wide association study was conducted using genome data of 199 samples collected from 14 countries that identified nine genetic loci associated with this phenotype and investigated genes in these regions that may contribute to the phenotype overall. Identifying the genetic factors may help better understand the biology and pathways that lead to the environmental effects on skin separation and improve commercial date production. In conclusion, our key finding is that both environmental and genetic factors contribute to skin separation variation, and improvements in environmental factors alone cannot overcome the extreme level of variation observed in some cultivars.

## 1. INTRODUCTION

Fruit quality is an essential factor for the market value in commercial agriculture. Biochemical composition and physical features are key parameters in determining quality, and the appearance of the fruit is a significant factor for consumers (Alam et al., 2023; Barrett et al., 2010; Mikulic-Petkovsek et al., 2021). The visual appeal of date palm fruits (dates) is adversely affected by exocarp separation from the mesocarp (commonly called skin separation or puffiness) and microcracks, which usually occur during the ripening stage (Alsmairat et al., 2023; Khadivi-Khub, 2014; Santos et al., 2023). Every year, growers experience significant financial loss because of fruit skin appearance (Alsmairat et al., 2023; Lobo et al., 2013). Skin separation phenomena affects the shelf-life of dates and customers’ purchasing choices (Lara et al., 2019). Understanding and addressing this phenotype in commercially important fruits, including dates, is a significant concern (Fernández-Muñoz et al., 2022; Lustig et al., 2014).

Date palm (Phoenix dactylifera L.) is an economically important crop in the Arabian Peninsula, North Africa and Pakistan, and plays a significant role in the economic development of these regions (Al-Shahib & Marshall, 2003; Chao & Krueger, 2007). Thousands of date palm varieties grow in hot, arid habitats worldwide; each variety exhibits a wide range of fruit characteristics, including sugar content, moisture, size and colour (Ahmed et al., 2016; Elhoumaizi et al., 2002; Younuskunju et al., 2023). Dates are a rich source of sugar, phenolic antioxidants, fiber, and proteins, making them economically significant worldwide (Al-Shahib & Marshall, 2003; Ghnimi et al., 2017). The uniformity of color and size, sugar content, and the absence of visual defects are some of the criteria for grading the quality of dates for marketing (Ahmed et al., 2016; Chao & Krueger, 2007; Elhoumaizi et al., 2002; Zaid & Arias-Jimenez, 1999). It is hypothesized that many of these these economically significant phenotypes are linked to genetic features and multiple recent studies have investigated this (Al-Dous et al., 2011; Hazzouri et al., 2019; Malek et al., 2020; Mathew et al., 2015; Younuskunju et al., 2023) including our own association study showing gentic control of dry fruit color (Tamar stage) (Younuskunju et al., 2023).

To take advantage of the long shelf life of dates requires the maintenance of undamaged skin throughout pre and post-harvest periods (Ahmed et al., 2021; Lara et al., 2019). The fruit skin (exocarp) is made up of composites of the cuticle, epidermis, and hypodermis and is considered an essential element in flesh fruits (Fernández-Muñoz et al., 2022; Gophen, 2014; Kayesh et al., 2013). It provides mechanical protection from biotic and abiotic stresses as well as contributes to the visual appearance (Ginzberg & Stern, 2016; Khadivi-Khub, 2014). Skin separation phenomenon does not occur in all date varieties and is mainly observed in economically significant varieties like Barhi, Sagai, Sukari, Khalas, Kheneizi, and Medjool (Alsmairat et al., 2023; Ghazzawy et al., 2023; Gophen, 2014; Lustig et al., 2014). Many studies have focused on understanding the factors causing this phenotype variation, ranging from microclimatic and nutritional aspects to mechanical characteristics of the date’s cell wall (Alsmairat et al., 2023; Ghazzawy et al., 2023; Gophen, 2014; Lobo et al., 2013; Lustig et al., 2014). Physio-chemical study of Sukari dates showed that environmental conditions, irrigation, and method of fertilisation are factors that can improve this and other fruit traits (Ghazzawy et al., 2023). Variations in the mechanical behaviour of different date cultivars, such as Dayri (no skin separation) and Barhi (extreme variation of skin separation), have been associated with skin separation phenomena (Gophen, 2014). Furthermore, the study of Medjool dates showed that environmental factors and cyclic stresses of turgor pressure fluctuations can also influence the traits (Lustig et al., 2014). Another study suggests that climate factors are not the only contribution to this phenotype (Alsmairat et al., 2023) by showing that the percentage of sclereid cells was significantly higher in the skin-separated fruit than in the normal one.

As we have observed, most previous studies examined the influence of environmental factors on skin separation in dates. However, we hypotheiszed that genetic factors may also play a part in determining which date cultivars are most affected by this phenotype. To our knowledge, no studies have been conducted to date to understand the influence of genetic factors on this disorder, likely because the impact of the environment is clear within a specific cultivar. Research on bell peppers has shown that genetic variation may impact skin separation (Kline et al., 2011; Wyenandt et al., 2017). The dataset used in this study is extensively diverse in origin and variety (Stephan et al., 2018) which is critical in understanding genetic association within the context of a phenotype also affected by environment. Importantly, because we collected the same cultivars from multiple environmental locations, our unique dataset could help distinguish the range environmental effect versus genetics on the skin separation phenotype.

## 2. MATERIAL AND METHODS

### Phenotypic data

Photographs of dry fruits of each cultivar were captured using a digital camera. Each cultivar had 5 to 11 fruits as representatives (Supporting Information S1: Figure 1). We manually assessed multiple images of each cultivar and scored the skin separation variation. The score was rated from 0 to 10 based on skin defects. Fruit with no defects was rated with a score of 0, and 10 was the maximum score for complete defects. A total of 1637 images belonging to 171 cultivars were manually assessed, and the skin defect rate was scored for each image. Outliers were removed from the raw dataset using the Z-score method ( used ± 2 standard deviations). The average skin separation rate was calculated using the outlier-removed raw data for each sample. We then performed a Boxcox transformation on the average score to reduce skewness. Box cox-transformed data were used as phenotypic data for the genome-wide association study.

**Figure 1:**
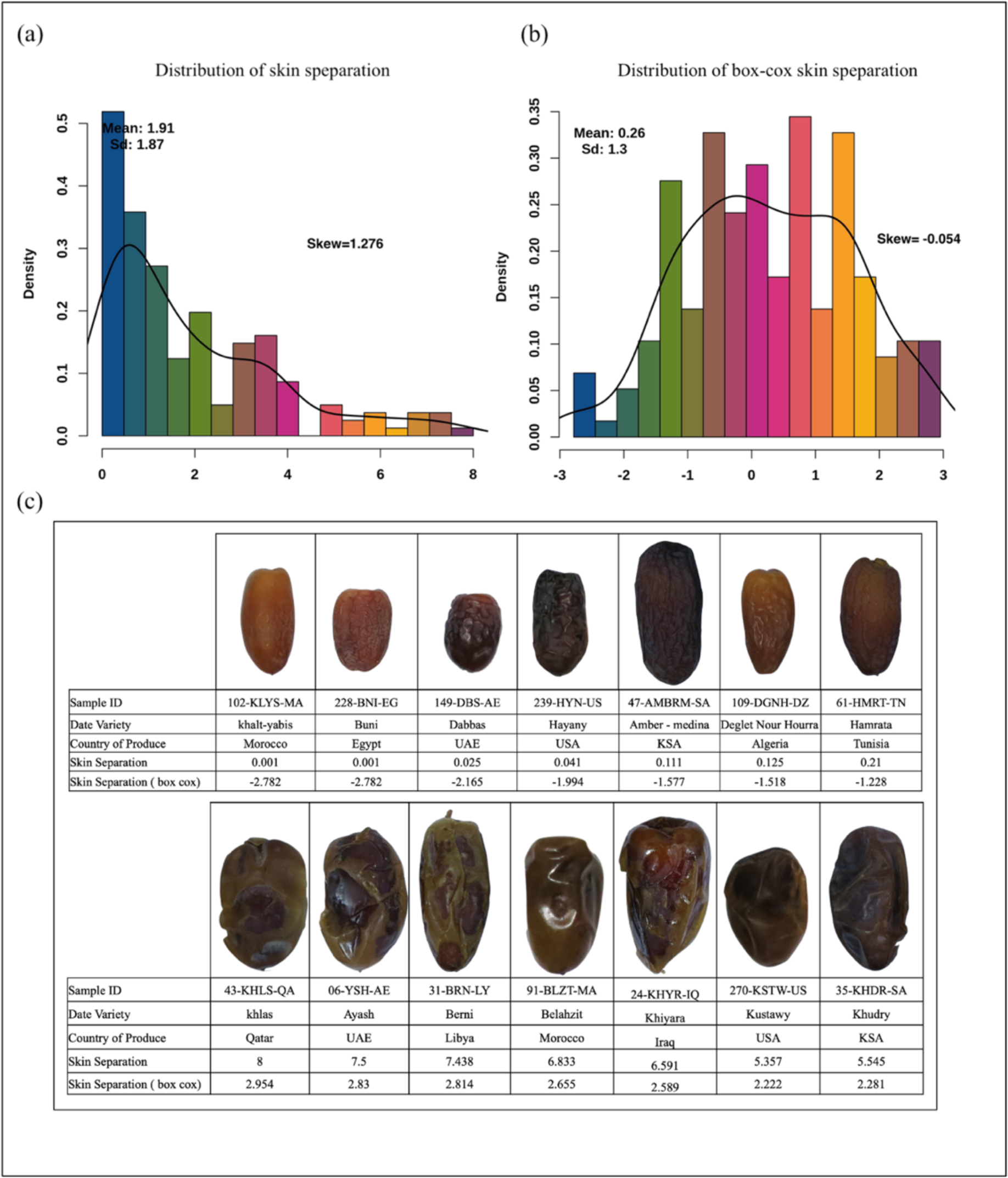
Distribution and comparison analysis of fruit skin separation phenotype values. We manually assessed multiple fruit images of each cultivar and scored skin separation ranging from 0 to 10. The distribution analysis showed that the phenotype data were positively skewed (right) with a skewness of 1.27, so we performed a Box-Cox transformation to reduce the skewness of the phenotypic data. (**a**) distribution analysis plot of raw phenotype values. (**b**) distribution plot of box-cox transformed phenotype values. (**c**) representative image of 7 date cultivars with the lowest and highest skin separation scores.

### Genome sequencing and SNP calling

We used a genome dataset of date samples from our previous association study of fruit colour (Younuskunju et al., 2023). Sequencing libraries were constructed from total DNA extracted from fruits, and whole genome libraries were sequenced using Illumina 2500/4000 instruments. For a more detailed description of the sequencing data, please refer to Mathew et al and Thareja et al study (Mathew et al., 2015; Thareja et al., 2018). The quality control processing of samples (QC), raw reads, genome alignment, and SNP calling was carried out in accordance with our previous association study on Tamar stage date fruit color (Younuskunju et al., 2023). SNPs were marked as missing if DP < 10 and filtered with the following parameters using VCftools (V0.1.16), genotype call rate 80%, minor allele frequency 0.01, and hardy-Weinberg equilibrium 1x10-6.

### Genetic similarity and phenotypic variation analysis

Genetic relatedness between the samples (kinship coefficients) was measured using Plink software (v1.9) (Purcell et al., 2007) with the ’ make-king table ’ option. The resulting data were filtered using a kinship score greater than or equal to 0.354 to find the genetically similar samples. Genetically similar samples were grouped based on pairwise kinship scores and considered for the phenotype variation analysis of genetically identical and dissimilar cultivars if a group had at least three or more samples from different regions. Skin separation variation analysis of samples within a group was performed to assess phenotypic differences between the genetically similar cultivars grown in different regions and environments. To compare the phenotypic variation between genetically different cultivars, we assessed the differences among samples from different groups.

### Genome-wide association study

Genome-wide analysis was performed (GWAS) using the FarmCPU method (Wang & Zhang, 2021) implemented in the GAPIT (v3) R package (Wang & Zhang, 2021). LD pruning was performed on the QC-filtered SNP dataset using the Plink software (--indep-pairwise 500 50 0.99) to improve the computational efficiency of the GWAS method (Purcell et al., 2007). A kinship matrix and four principal components (PCA) were used as covariates in the GWAS to correct the population structure. Both the PCA and kinship matrix scores were calculated from the LD-pruned SNPs using the GAPIT R package. The VanRaden algorithm in the GAPIT R package was used to measure the kinship matrix. A list of significant SNPs associated with phenotype was identified using a cutoff value of FDR-adjusted p-values of 0.05 ( 5 %). Manhatten and QQ plots were generated from association results using the CMPlot R package (Yin et al., 2021).

### Structural variation and RNA seq analysis candidate gene

Regions spanning 100 kb upstream and downstream of GWAS-identified significant SNPs were examined to determine potential candidate genes and variants. The gene sequences were obtained from these regions using the GFF3 annotation file of the PDK50 reference genome (PRJNA40349). Gene ontology analysis of the candidate gene sequences was conducted using Blast2Go software (Conesa & Götz, 2008). Gene functions were determined through literature review and Blast2Go results. All INDELs and SNPs from the regions were annotated using SNPEff software (Cingolani et al., 2012). RNA-seq expression analysis was carried out using the transcriptome data of kheneizi and Khalas cultivars from Hazzouri et al.’s study (Hazzouri et al., 2015). The data contains three or four replicates taken at different post-pollination days in two cultivars (45,75,105,120,135 days). Reads alignment and expression analysis was performed as described in our previous association study on fruit color (Younuskunju et al., 2023). Structural variation analysis was conducted as described in the association study on fruit color (2023).

## 3. RESULTS

### Phenotype data analysis: Skin separation rate

Manual analysis of the fruit images showed that the rate of skin separation varied from cultivar to cultivar (Figure 1). The averaged data set showed that 62 samples had a skin separation score greater than or equal to 2, while 109 samples had a score less than 2. The minimum score observed was 0, and the maximum score reached 8. The distribution analysis of phenotype indicated a positive skew (right) with a skewness value of 1.27 ( Figure 1a). To improve the data quality and reduce skewness for the association study, a box-cox transformation was carried out (Cieleń et al., 2023). This transformation process reduced the skewness of the phenotypic data to -0.054 (Figure 1b). During the transformation, the scores less than or equal to 0.99 were converted to negative values, and scores greater than or equal to 1 were converted to positive values (Figure 1c). This box-cox transformed dataset was used as the phenotypic data for the association study (Supporting Information S2).

### Genotyping and SNP calling

Quality control (QC) filters of the raw genotypic dataset produced 188 samples and 10,183,993 SNPs. These SNPs span over 18 linkage groups (LGs) and unplaced scaffolds in the reference genome. The genomes of the GWAS population showed that the correlation coefficient value (R2) decreased to half its maximum at 25.9 Kb (half LD decay value). For more detailed results of SNP calling and LD decay analysis, please refer to the study by Younuskunju et al (Younuskunju et al., 2023). LD pruning resulted in 3.541 million SNPs that were utilized in the association study.

### Genetic relatedness and phenotypic variation analysis

The genetic relatedness analysis resulted in the identification of sample pairs with kinship coefficient scores (Supporting Information S1: Figure 2). Samples with estimated kinship coefficient scores above 0.354 were considered genetically similar. Based on their kinship coefficient scores, these genetically similar samples were sorted into 23 sample groups. To compare the skin separation differences among genetically similar grown in different locations and environments, we chose five sample groups out of the 23, each containing at least three or more samples from different regions (Supporting Information S1: Table 1). These groups are named Deglet Nour, Medjoul, Mabroom, Safawi, and Sagai. The analysis of phenotype variation was carried out within each group separately, and then averages were compared between groups (Figures 2 and 3). The samples in the Deglet Nour group exhibited scores ranging from 0.12 to 1.1,Medjoul from 1 to 2, Mabroom from 2 to 3.63, Safwai from 1.49 to 2.72, and Sagai from 3.9 to 5.04, and so on (Figure 2). The phenotype comparison between different cultivar groups showed higher phenotypic differences than did the intra-cultivar comparison (Figure 3). The Deglet Nour group showed an average score of 0.58, Medjoul an average of 1.76, Mabroom an average of 2.75, Safawi an average of 2.06, and Sagai an average of 4.46. The comparison of the standard deviation of the average score between genetically similar and dissimilar cultivars (Degelt Nour, Medjoul & Sagai) showed considerable differences in phenotype [Figure 3b]. Samples within the Deglet Nour group show a standard deviation of 0.4, the Medjoul group 0.41, and the Sagai group 0.6. The standard deviation between the Degelt Nour and Medjoul group was 1.01, the Medjoul and Sagai group was 1.72, and the Deglet Nour and Sagai group was 2.72. The comparison of the Deglet Nour group to the Sagai group showed the most extreme difference in skin separation between groups.

**Figure 2:**
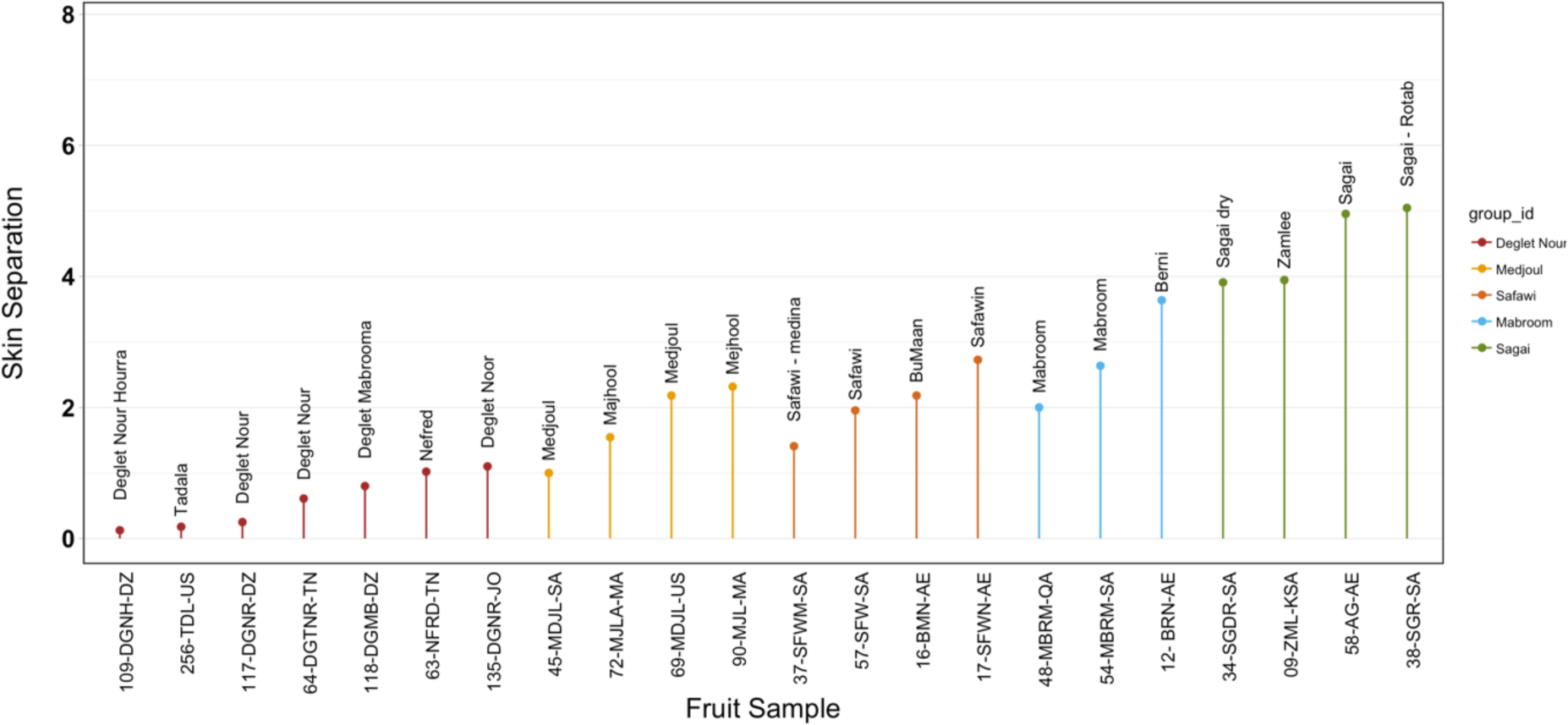
Comparison analysis of the skin separation variation of genetically similar fruit cultivars grown in different regions and environments. Genetically similar cultivars were marked with a separate colour code. The X-axis represents samples from multiple cultivars, and the Y-axis represents the skin separation rate.

**Figure 3:**
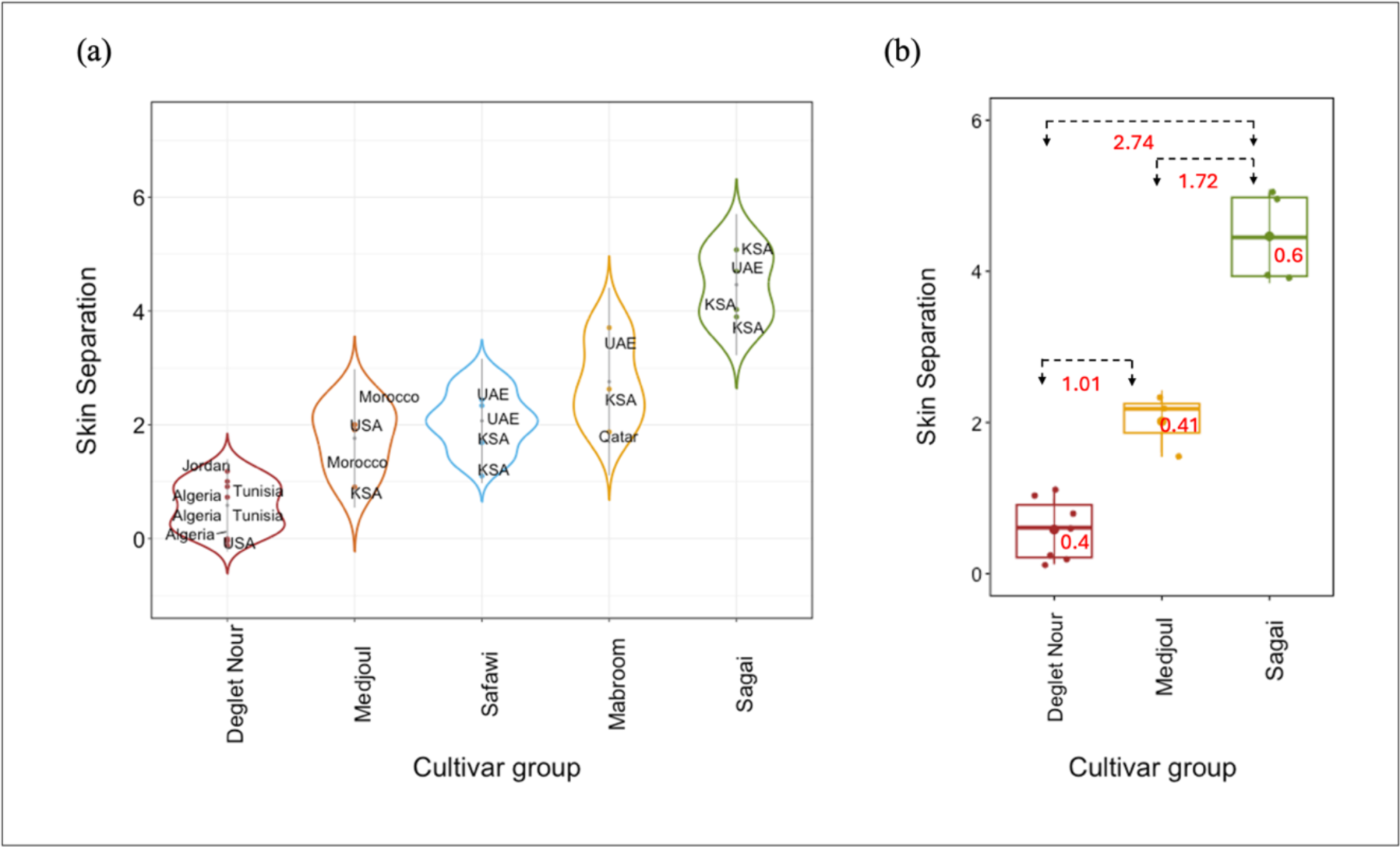
Comparison analysis of fruit skin separation of genetically different fruit cultivars. (**a**): Skin separation comparison of five genetically different cultivars. Each point represents a sample and is marked as the country of origin. (**b**): boxplot distribution and standard deviation analysis of skin separation in three genetically different cultivar groups. The standard deviation of skin separation scores for fruit samples within each cultivar group and between the groups is marked in red color. The X-axis represents the cultivar group, and the Y-axis represents the skin separation rate.

**Table 1:** List of significant SNPs associated with skin separation phenotype from association study (GWAS). A false-discovery rate (FDR) adjusted p-value was used as a cut-off for identifying significant SNPs associated with the phenotype.

| SNP | LD ID | MAF | P.value | FDR P.Value |
| --- | --- | --- | --- | --- |
| LG8s1288981 | LG8 | 0.14 | 5.71E-16 | 2.02E-09 |
| LG3s2983485 | LG3 | 0.21 | 2.26E-14 | 4.00E-08 |
| LG6s6604213 | LG6 | 0.07 | 1.34E-11 | 1.58E-05 |
| LG6s3747705 | LG6 | 0.05 | 1.11E-10 | 9.64E-05 |
| LG18s8763935 | LG18 | 0.46 | 1.36E-10 | 9.64E-05 |
| LG4s35891773 | LG4 | 0.05 | 1.46E-09 | 0.00086216 |
| LG10s3647074 | LG10 | 0.35 | 2.01E-08 | 0.01017247 |
| LG2s54442820 | LG2 | 0.09 | 3.57E-08 | 0.01581493 |
| LG18s848875 | LG18 | 0.22 | 5.76E-08 | 0.02267 |

### Association of fruit skin separation and Significant SNP’s genotypic effects

Genome-wide association using the box-cox transformed phenotypic data resulted in the discovery of several significant SNPs. The QQ plot of the association showed a lambda score of 1.02, indicating that the test statistics aligned with the expected distribution (Figure 4a). The FDR-adjusted p-value cutoff of 5% ( FDR < 0.05) identified nine SNPs that were significantly associated with the phenotype (Table 1; Figure 4b; Supporting Information S3). These SNPs were located in multiple LGs in the PDK50 reference genome. Among the identified significant SNPs, LG8s1288981, LG3s2983485, LG18s8763935, LG10s3647074, and LG4s35891773 SNPs showed significant Wilcoxon test p-values (Figure 5, Supporting Information S1: Figure 3). Analysis of the 3 possible genotypes at the associated SNPs showed the effect of each allele (Figure 5). Further validation was provided by the visual examination of the phenotype when separated by the genotypes of SNP LG18s8763935 (Figure 6). It was observed that the fruit exhibited a higher level of skin separation rate when the sample was homozygous for the reference allele (REF) at SNP LG18s8763935, whereas the fruit showed a lower rate when the sample was homozygous for the alternative allele (ALT) at SNP LG18s8763935.

**Figure 4:**
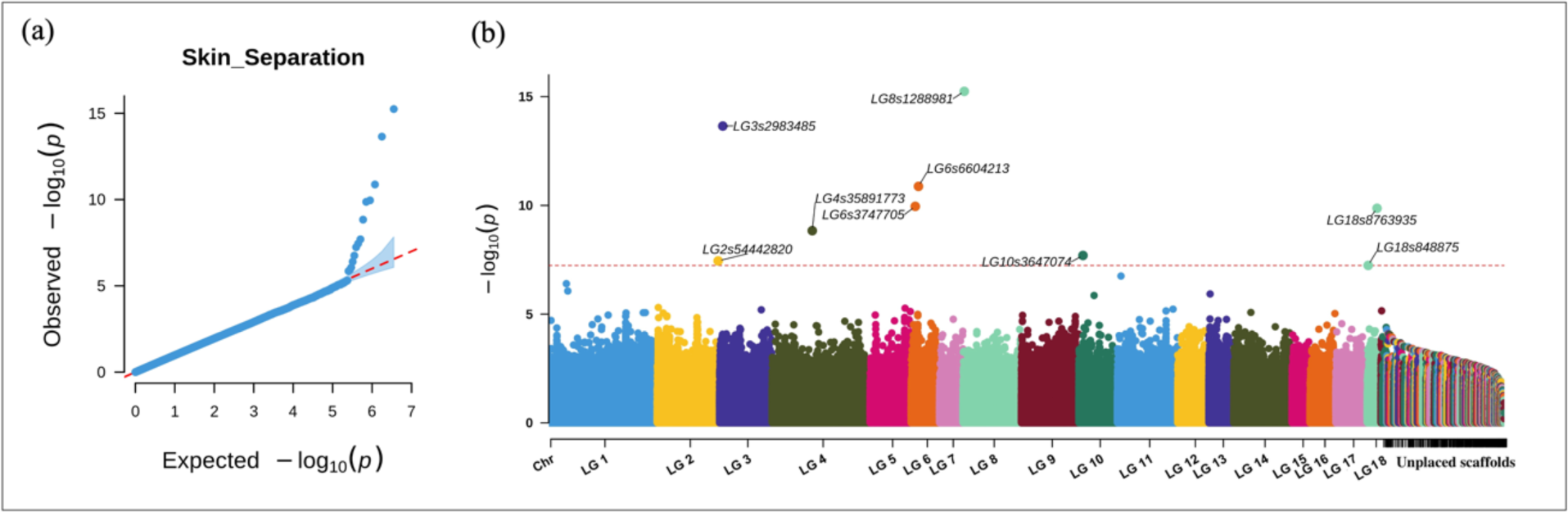
Genome-wide association study (GWAS) analysis of skin separation phenotype using the LD pruned SNP set of 3.541 million SNPs.(**b, c**) QQ plot and Manhattan plot using the LD pruned SNP set for all Linkage group and unplaced scaffolds.

**Figure 5:**
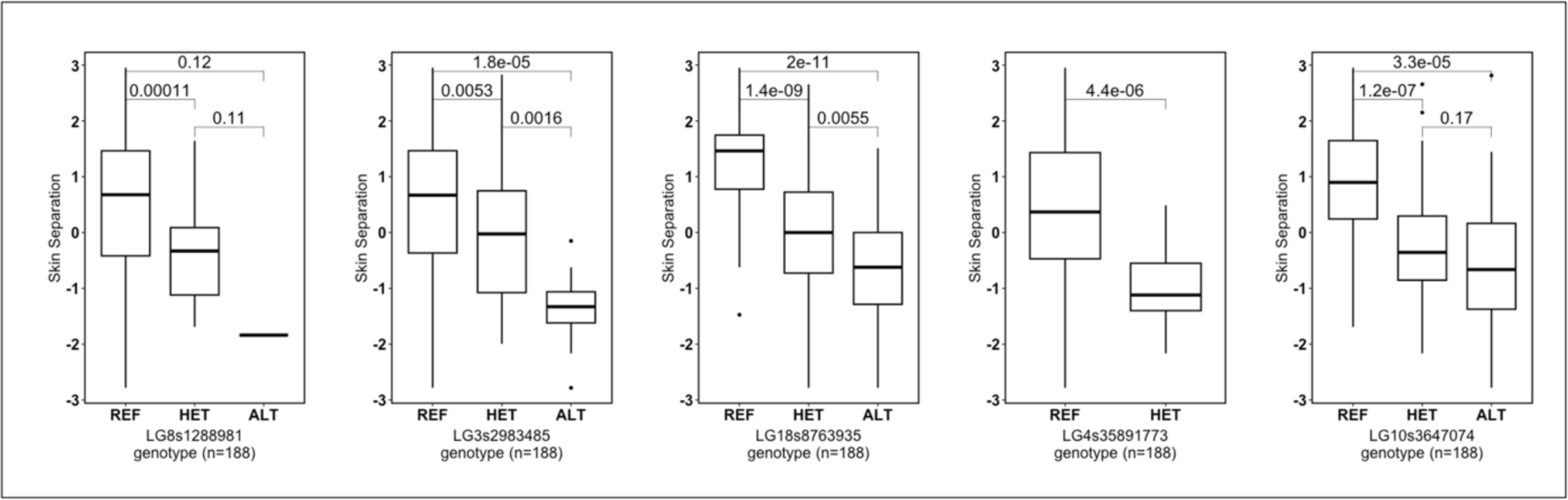
Boxplot distribution analysis of skin separation phenotypes by genotypes of a list of significant SNPs from this association study (GWAS). These five SNPs have significant Wilcoxon test p-values among the list of GWAS identified SNPs. The *X*-axis represents the SNPs’ genotypes, and the *Y*-axis represents the phenotypic value (box-cox transformed phenotype).

**Figure 6:**
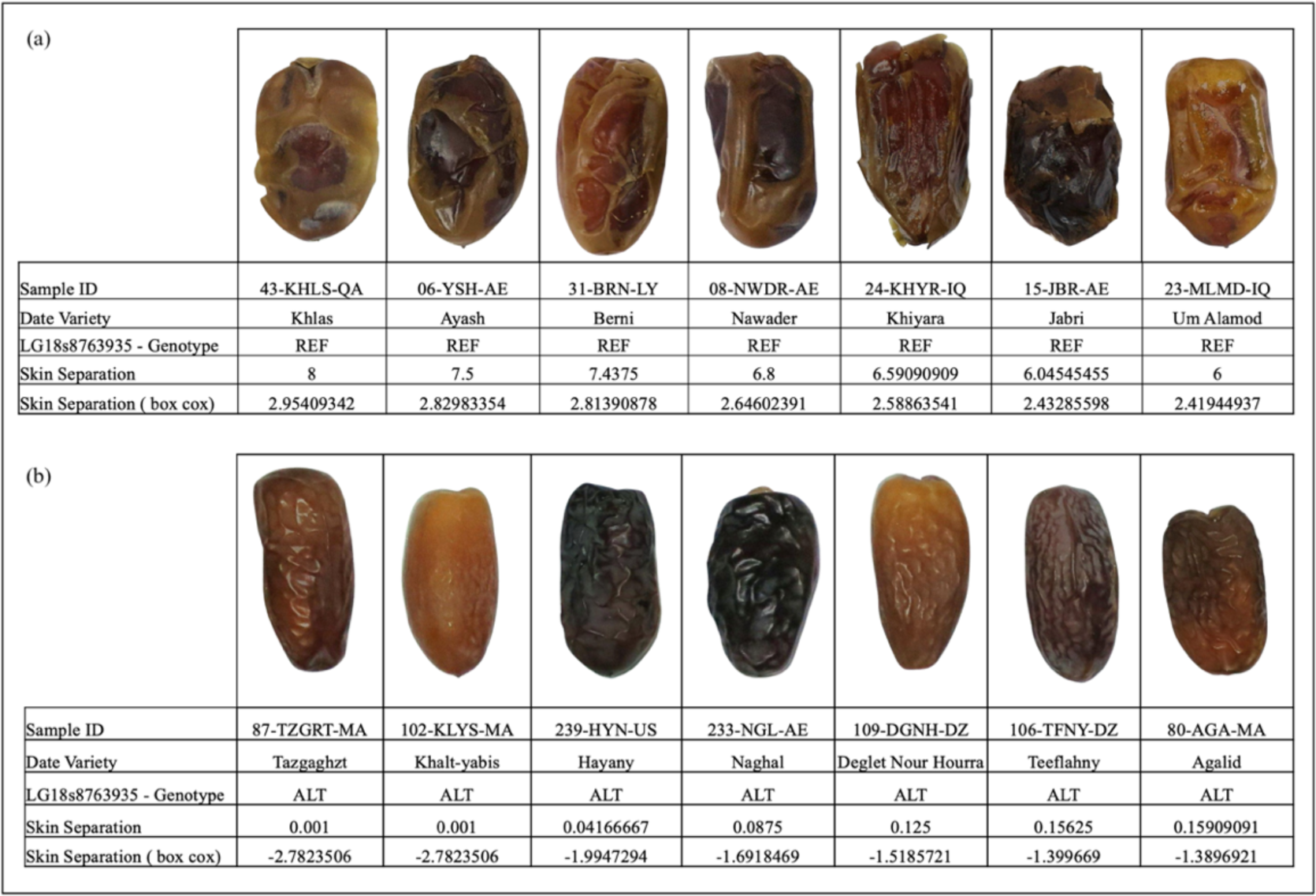
Differences in fruit skin separation when samples were categorised by the genotypes of LG18s8763935 SNP. Fruits were grouped together which are homozygous for the **(a)** RFE or **(b)** ALT allele of SNP LG18s9876335. Results show that fruit skin separation is at an extreme level when the sample is homozygous for the REF allele and at a very low level or none when the sample is homozygous for the ALT allele at SNP LG18s8763935.

### Candidate gene and SNP annotation

A total of 150 genes were identified across the potential regions (Supporting Information S4). The results of literature searches and Blast2Go showed that many of these genes are involved in lignin synthesis, plant-type cell wall loosening, cell wall organisation, and response to auxin, abscisic acid, and gibberellic acid (Table 2). Gene expression analysis using RNA-seq data from Khalas and Kheneizi varieties showed that many of these genes were indeed expressed during various stages of fruit development (post-pollination days) (Figure 7). Expansin, Cellulose synthase-like protein, Myb transcription factor, Ras-related protein Rab-8A, C2 and GRAM domain-containing protein, Soluble inorganic pyrophosphatase, Transport inhibitor response 1- like protein genes were expressed during the early stages of development (dpp 45-70) and the Proteasome subunit beta type, DOF zinc finger protein 1 were expressed during the later stages (dpp 105- 135). The SNPEff annotated SNPs and INDELs from the candidate genes with an LD R2 value >=0.6 to the significantly associated SNPs showed many putative high and moderate effects to encoded proteins (Supporting Information S4). Structural variation (SV) analysis was performed in all potential regions of the significantly associated SNPs. However, no significant SVs were found in the potential regions of the significantly associated SNPs.

**Figure 7:**
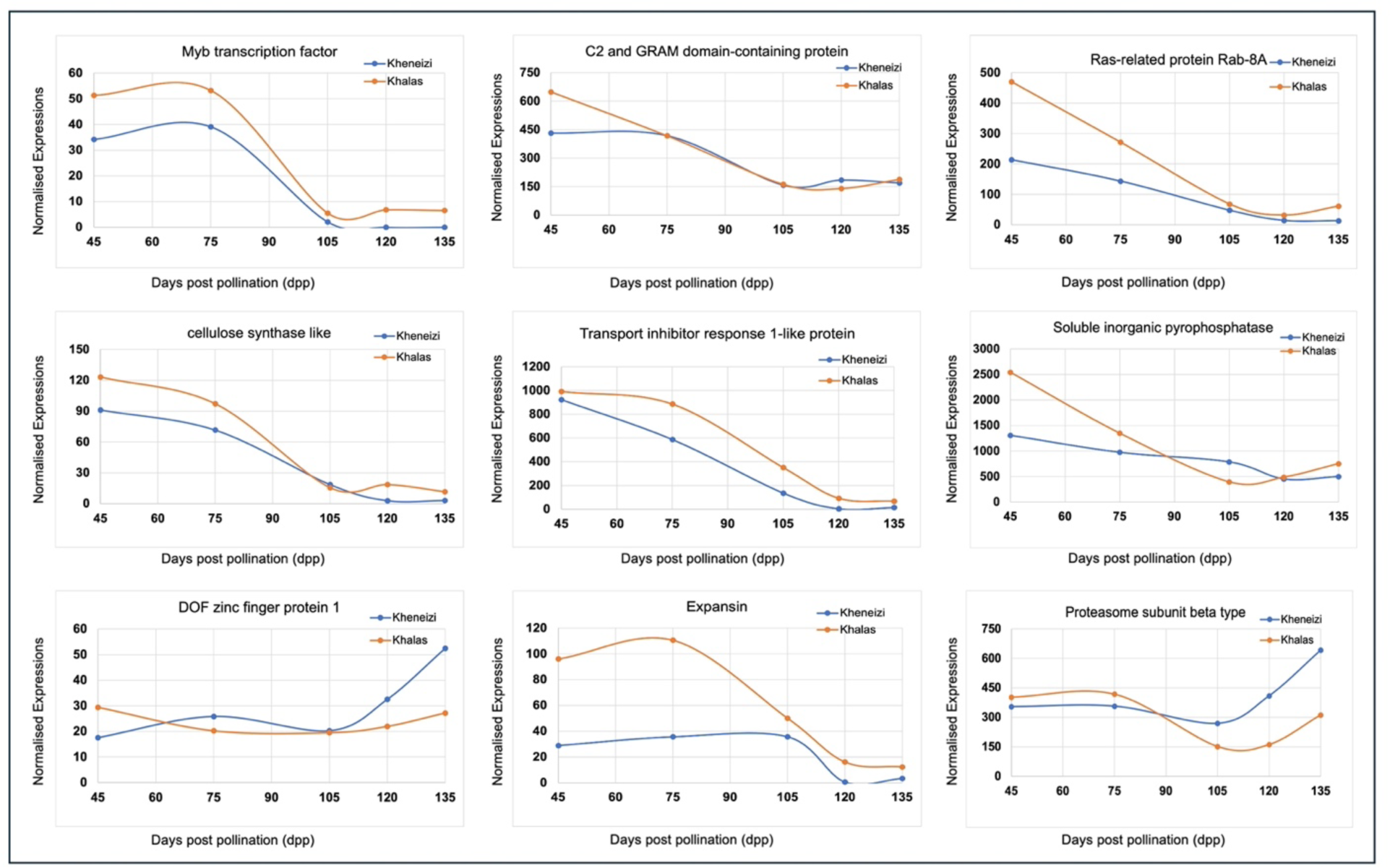
RNA seq analysis of expression of genes identified in the potential region around identified SNPs. The gene expression analysis was carried out across three or four replicates of fruit development stages of two varieties, namely Kheneizi (red) and Khalas (yellow). Each point on the X-axis represents the development stage (post-pollination date), while the Y-axis displays the mean normalised expression read count across three or more replicates.

**Table 2:** List of genes detected around the 100kb region of significant SNPs from association study (GWAS) result associated with the skin separation phenotype. Genes were selected if they have a putatively significant role in fruit growth regulation, lignin synthesis, and plant-type cell wall development.

| Tag SNP | Protein Name | Gene function |
| --- | --- | --- |
| LG18s8763935 | Proteasome subunit beta type | response to water deprivation; response to ethylene; response to abscisic acid; brassinosteroid mediated signaling pathway; positive regulation of auxin mediated signaling pathway; |
| LG3s2983485 | basic helix-loop-helix (bHLH) DNA-binding superfamily protein | response to gibberellin; cellular response to water deprivation; cellular response to abscisic acid stimulus; |
| LG3s2983485 | Soluble inorganic pyrophosphatase | cell wall thickening; cell wall pectin metabolic process; |
| LG3s2983485 | Ras-related protein Rab-8A | cell wall biogenesis |
| LG4s35891773 | C2 and GRAM domain-containing protein | positive regulation of abscisic acid-activated signaling pathway |
| LG6s6604213 | Expansin | response to gibberellin; unidimensional cell growth; plant-type cell wall loosening; structural constituent of cell wall; |
| LG6s6604213 | Myb transcription factor | response to water deprivation; response to ethylene; response to abscisic acid; organ boundary specification between lateral organs and the meristem; positive regulation of auxin mediated signaling pathway; |
| LG6s6604213 | DOF zinc finger protein 1 | response to auxin; response to salicylic acid; cell wall modification; positive regulation of cell cycle; |
| LG6s6604213 | cellulose synthase like | cell wall organization; |
| LG6s6604213 | cellulose synthase like | cell wall organization |
| LG6s6604213 | Cellulose synthase-like protein | cell wall organization |
| LG8s1288981 | Cyclin-D4-1 | cell division |
| LG8s1288981 | Cyclin-D3-1 | response to cytokinin; response to sucrose; guard mother cell differentiation; cell division; |
| LG8s1288981 | DNA-directed RNA polymerase D subunit 1-like protein | regulation of cell division; |
| LG8s1288981 | Transport inhibitor response 1-like protein | auxin-activated signaling pathway; response to jasmonic acid; |

## 4. DISCUSSION

Previous studies on date skin separation phenomena have demonstrated strong evidence that environmental factors play a significant role in the phenotype (Alsmairat et al., 2023; Bitar, 2021; Ghazzawy et al., 2023; Gophen, 2014; Lustig et al., 2014). Determining the genetic association with a phenotype in the presence of environmental factors presents a considerable challenge (Aschard et al., 2012; Wojcik et al., 2019). The study requires vast and diverse sample data sets from various environments. In this study, we used a unique dataset to determine the effect of genetic factors on skin separation phenomena with environmental backgrounds. We collected a variety of genetically similar cultivars from different environments and locations. The samples, which include the same cultivars from multiple environmental and global locations, enable us to determine the extent to which the environment versus genetics influence the trait. The analysis showed a range of phenotype variations in genetically similar (intra-cultivar analysis) and between cultivars (inter-cultivar analysis). The central question is whether this phenotypic difference is solely influenced by environmental factors or if the contribution of genotypic factors, in addition to environmental factors, leads to the variation.

The analysis of genetically similar cultivars grown in different regions showed the range of skin separation between the samples of the same cultivar (Figure 2) based on environment. That is, the phenotypic range in these cultivars is due to differences in factors such as watering, post-harvest treatment and other abiotic affects. We observe significant differences highlighting the importance of environmental factors on skin separation. However, importantly, the analysis conducted between genetically different cultivars demonstrated an extreme level of variation compared to the variation in genetically similar cultivars grown in distinct regions and environments [Figure 2, Figure 3]. The score varies from an average of 0.5 to 4.46 [Figure 3]. The variation within a cultivar is less than that observed between the most affected cultivars. The standard deviation between genetically different cultivars is higher than the standard deviation between the samples of genetically similar cultivars present within the group (Figure 3b). These extreme differences between cultivar groups are likely due to genetics. That is, genetic and environmental factors contribute to the differences in skin separation phenotype, but the genetic factor is the strongest contributor to extreme differences.

Based on these observations, we conducted an association (GWAS) study on date varieties, including genetically similar cultivars grown in different locations, to further investigate the genetic influence on the phenotype. The GWAS results showed nine markers associated with the phenotype. The analysis of phenotypes by genotypes demonstrated that the genotypes of these SNPs are significantly associated with phenotypic variation (Figure 5, Supporting Information S1: Figure 3). Studies have shown that changes in the biochemical properties of exocarps during fruit ripening can lead to skin separation and microcracking (Lara et al., 2019; Petit et al., 2014). Identifying the genes responsible for the development and modification of cell walls and cuticular membranes will lead to a better understanding of genetic factor’s contribution to these skin disorders. Our study identifies several key genes involved in cell wall development and modification in the potential candidate regions of significant markers [Table 2]. The Expansin gene is present in the candidate region of LG6s6604213 SNP. This gene plays a crucial role in cell wall loosening and weakening during cell expansion (Brummell et al., 1999; Cosgrove, 2000).

Cosgrove’s (2000) study showed that plant cells produce expansin protein during growth, which unlocks the polysaccharide wall network and allows turgor-induced cell walls to loosen (Cosgrove, 2000). The candidate regions from LG3s2983485, LG4s35891773, LG8s1288981, and LG18s8763935 SNPs contain many genes that have functional responses to growth regulator genes such as Auxin, Gibberellic acid (GA), and abscisic acid. Studies in apples and litchi have shown that these growth regulators have influenced skin cracks during fruit development (Joshi et al., 2018; Khadivi-Khub, 2014). Studies on apples show that epidermal density is associated with the cracking resistance of the fruit (Eccher & Hajnajari, 2005; Joshi et al., 2018). These studies suggested that the level of GA could increase the epidermal cell density. The candidate region of the LG3s2983485 SNP contains the pyrophosphatase gene, which is involved in cell wall thickening and the metabolic process of cell wall pectin (Li et al., 2014).

Our study findings help reveal the various contributions of environmental and genetic factors to skin-separation phenotypic variation in dates. We confirm that environmental factors likely modify skin separation given intra-cultivar variation. However, in some cultivars like Sagai, the genetic factors are so signficant that environmental improvement may only result in minor effects improvements. Our study contributes to the understanding of the influence of environmental and genetic factors on skin separation in the most popular date palm cultivars. This knowledge will benefit growers in selecting and developing fruit varieties with reduced skin separation in breeding programs.

## Data archiving statement

Table 1 includes the SNP association results of Date palm skin separation phenotypes on the date palm reference genome. Gene annotation from the associated linkage group is included in Table 2.

## Author contributions

SY designed the study, analysed data, and wrote the manuscript; LSM conducted genome sequencing; YAM directed library construction and sequencing; JAM envisioned the project, conducted bioinformatics analysis and wrote the manuscript; KS designed the study and helped Write the manuscript. KFXM designed the study and helped write it.

## Compliance with Ethical Standards

Human and animal subjects were not included in this study.The authors declare not competing interests.

## Acknowledgements

We thank Robert Krueger of the USDA, Sean Lahmeyer of Huntington Gardens and Diego Rivera of the University of Murcia for contributing date palm, Phoenix and palm species.

