## Supplemental Table 1 and Figure for "The genetics of fruit skin separation in date palm"

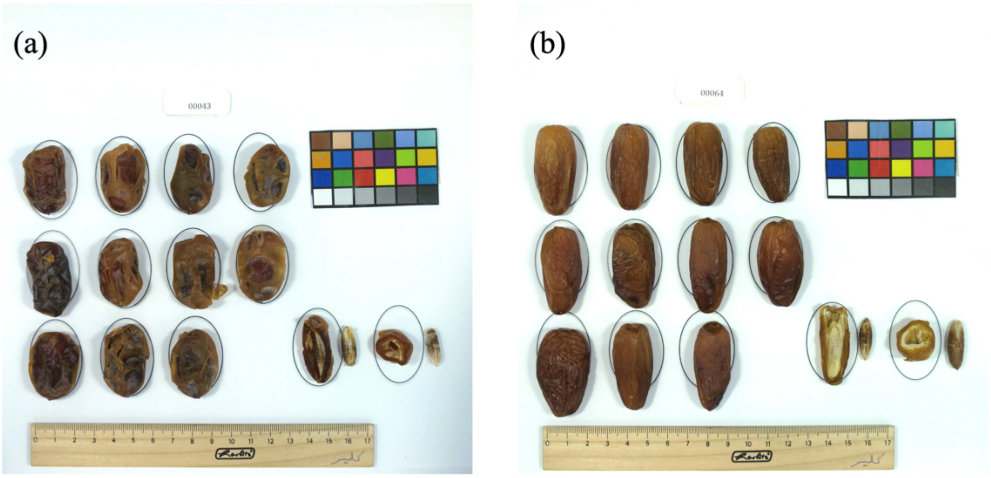


**Figure 1**: Fruits Photograph used for scoring the fruit's skin separation rate. The score was rated from 0 to 10 based on skin separation. Fruits with no skin separation were rated with a score of 0, and 10 was the maximum score for fruit with full skin separation. (a, b)  Fruits Photographs of cultivars used for phenotype measurement.

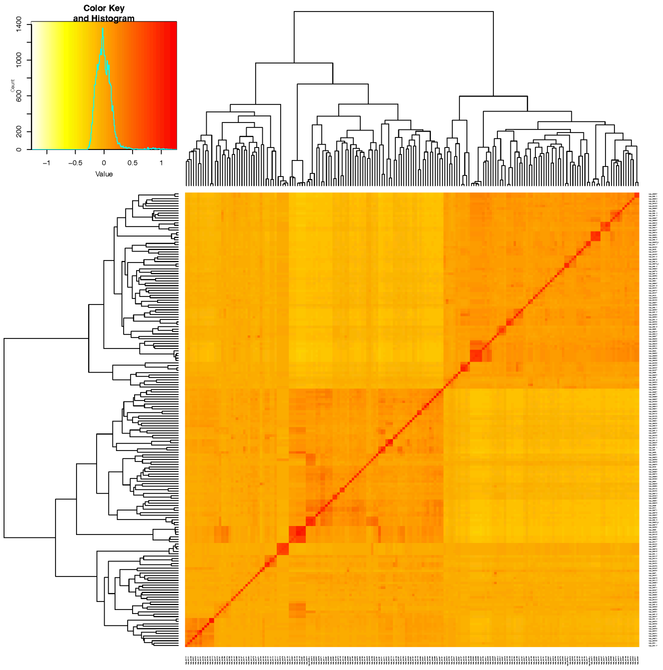


**Figure 2**: Heatmap of pairwise kinship matrix values, based on all LD pruned SNPs on 188 samples, according to the VanRaden algorithm. The color histogram shows the distribution of coefficients of co-ancestry, and the stronger red color indicates that individuals are more related to each other.

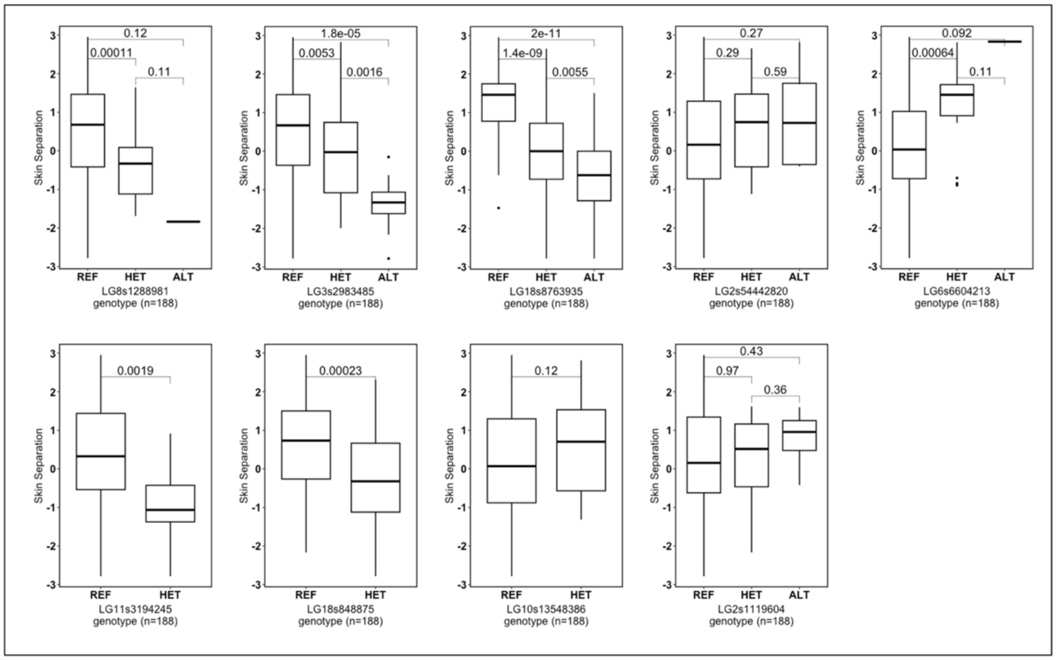


Figure 3: Boxplot distribution analysis of skin separation phenotypes by genotypes of a list of significant SNPs from this association study (GWAS). The *X*-axis represents the SNPs' genotypes, and the *Y*-axis represents the phenotypic value (box-cox transformed phenotype).

**Table 1:** List of samples and cultivar groups used for skin separation variation analysis in genetically similar and dissimilar cultivars. Cultivar groups were considered for analysis if a group had at least three or more samples from different regions, ensuring a diverse and representative sample.

| Sample ID | Date variety | Country of produce | Skin separation score | Cultivar group |
|---|---|---|---|---|
| 109-DGNH-DZ | Deglet Nour Hourra | Algeria | 0.13 | Deglet Nour |
| 256-TDL-US | Tadala | USA | 0.18 | Deglet Nour |
| 117-DGNR-DZ | Deglet Nour | Algeria | 0.25 | Deglet Nour |
| 64-DGTNR-TN | Deglet Nour | Tunisia | 0.61 | Deglet Nour |
| 118-DGMB-DZ | Deglet Mabrooma | Algeria | 0.80 | Deglet Nour |
| 63-NFRD-TN | Nefred | Tunisia | 1.02 | Deglet Nour |
| 135-DGNR-JO | Deglet Noor | Jordan | 1.10 | Deglet Nour |
| 37-SFWM-SA | Safawi - medina | KSA | 1.41 | Safawi |
| 57-SFW-SA | Safawi | KSA | 1.95 | Safawi |
| 16-BMN-AE | BuMaan | UAE | 2.18 | Safawi |
| 17-SFWN-AE | Safawin | UAE | 2.73 | Safawi |
| 45-MDJL-SA | Medjoul | KSA | 1.00 | Medjoul |
| 72-MJLA-MA | Majhool | Morocco | 1.55 | Medjoul |
| 69-MDJL-US | Medjoul | USA | 2.18 | Medjoul |

| | | | | |
|---|---|---|---|---|
| 90-MJL-MA | Mejhool | Morocco | 2.32 | Medjoul |
| 48-MBRM-QA | Mabroom | Qatar | 2.00 | Mabroom |
| 54-MBRM-SA | Mabroom | KSA | 2.64 | Mabroom |
| 12- BRN-AE | Berni | UAE | 3.64 | Mabroom |
| 34-SGDR-SA | Sagai dry | KSA | 3.91 | Sagai |
| 09-ZML-KSA | Zamlee | KSA | 3.94 | Sagai |
| 58-AG-AE | Sagai | UAE | 4.95 | Sagai |
| 38-SGR-SA | Sagai - Rotab | KSA | 5.05 | Sagai |
